# Probing the Role of Membrane in Neutralizing Activity of Antibodies Against Influenza Virus

**DOI:** 10.1101/2025.02.11.637756

**Authors:** Defne G. Ozgulbas, Timothy J. C. Tan, Po-Chao Wen, Qi Wen Teo, Huibin Lv, Zhaleh Ghaemi, Martin Frank, Nicholas C. Wu, Emad Tajkhorshid

## Abstract

Influenza poses a major health issue globally. Neutralizing antibodies targeting the highly conserved stem region of hemagglutinin (HA) of the influenza virus provide considerable protection against the infection. Using an array of advanced simulation technologies, we developed a high-resolution structural model of full-length, Fab-bound HA in a native viral membrane to characterize direct membrane interactions that govern the efficacy of the antibody. We reveal functionally important residues beyond the antibody’s complementary-determining regions that contribute to its membrane binding. Mutagenesis experiments and infectivity assays confirm that deactivating the membrane-binding residues of the antibody decreases its neutralization activity. Therefore, we propose that the association with the viral membrane plays a key role in the neutralization activity of these antibodies. Given the rapid evolution of the influenza virus, the developed model provides a structural framework for the rational design and development of more effective therapeutic antibodies.

## INTRODUCTION

Influenza virus remains responsible for millions of deaths annually due to respiratory illnesses caused by both its seasonal and occasional pandemics. The rapid evolution of the virus, driven by the antigenic drift of its surface glycoproteins, necessitates the annual administration of reformulated vaccines to protect against severe outcomes associated with its infection^1,2^. Typically, vaccines target the virus’ most abundant surface glycoprotein, hemagglutinin (HA), which continues to serve as the primary antigen for development of anti-influenza therapeutics and vaccines^3–5^.

HA is a trimer of identical subunits consisted of membrane-distal HA1 and membrane-proximal HA2 fragments, tethered through disulfide linkages^6^. The protein is vital for infection, playing a central role in two critical steps: binding of the virus to cell-surface sialic acid receptors, and subsequent fusion of the viral membrane with the host cell membrane^7,8^. These processes rely on HA large-scale conformational changes, which are primarily induced by pH variations^9^.

While antibodies can recognize HA and block receptor binding and/or membrane fusion to protect against influenza, most antibodies that bind HA at variable regions located within the membrane-distal HA1 provide only limited protection against a small number of related viral strains. As a result, broadly neutralizing antibodies (bnAbs)^10–14^ have been developed as a promising strategy for preventing viral infections more effectively. They target conserved regions of the HA protein, such as the membrane-proximal stem domain in HA2, thereby inhibiting the viral entry through membrane fusion^15,16^.

One specific human monoclonal bnAb against H1 strains, FISW84, has drawn significant attention due to its highly conserved epitope target that is shared with several other stalk-binding antibodies^12^. FISW84 binds near the junction between the ectodomain and the transmembrane (TM) domain of HA2^13,17,18^, which is also targeted by other broadly neutralizing antibodies that work against most H1 subtypes of influenza A viruses^12,17^. Notably, this binding position places FISW84 in close proximity to, actually overlapping with, the viral membrane, raising questions about the membrane’s potential involvement in the antibody’s binding and neutralization activity. In other viral infections, such as HIV, bnAbs targeting the membrane-proximal stem region, e.g., 4E10, 2F5, and 10E8, require initial association with the viral membrane to execute their neutralization effect^19,20^. Moreover, it has been shown that neutralization depends not only on the strong interaction between the antigen-binding fragment (Fab) of antibodies and viral glycoproteins but also on semi-specific interactions with membranes and specific binding to phospholipid head groups^21–26^.

While the mechanism of influenza antibodies binding to the stem region of HA and preventing its low-pH conformational change remains poorly understood, structural hints at possible interaction with the membrane hold significant implications for viral entry and for design of improved therapeutics. In this study, we adopt advanced computational modeling and *µ*s-scale molecular dynamics (MD) to construct and simulate a model for antibody-bound HA in the explicit presence of the viral membrane with the goal of unraveling the atomistic interactions between FISW84 and the membrane when complexed with HA. We first construct full models of HA starting from the cryo-EM structure of the A/duck/Alberta/35/76 (H1N1) viral strand (PDB ID: 6HJQ), the influenza antibody FISW84-Fab, and a realistic lipid bilayer that mimics the composition of the influenza viral membrane. Next, we employ all-atom MD simulations to explore the role of the viral membrane in stabilizing the Fab binding to HA and characterize the interfacial residues responsible for contacts with the membrane, which are likely to affect the binding of FISW84. Using these predicted sites, we then design and perform mutagenesis experiments and neutralization assays to validate their involvement in membrane binding and neutralization activity of the antibody. By comprehensively analyzing the conformational changes, stability, and interactions within the HA-Fab-membrane system, we provide a structural framework to guide for the development of potential therapeutic approaches in combating influenza.

## RESULTS AND DISCUSSION

In light of the crucial role of HA in influenza virus entry into the host cell and the growing interest in developing more effective antiviral strategies, the main objective of the present study is to characterize the unknown structure of the HA-Fab-membrane ternary complex. In the following sections, we first lay out our integrative modeling/simulation approach to construct a structural model for the HA-Fab complex anchored in a viral membrane and then describe MD simulations performed on the model to gain structural and dynamical insights into the role of the membrane in viral neutralization effect of the antibody. Mutagenesis experiments allowed us to validate key interactions between a number of non-paratope-binding residues in the antibody and the lipids of the membrane captured by the simulations. Additionally, we explored the impact of varying the number of bound Fab domains on the structure and dynamics of the complex with simulations.

### Construction of membrane-embedded, Fab-bound HA

To gain structural insight into the involvement of the membrane in binding and neutralization activity of stem antibodies, we first constructed a membrane-embedded, Fab-bound HA using a multi-step integrative approach. The cryo-EM structure available for the FISW84 Fab and HA complex^17^ lacks any representation for the viral membrane. Nevertheless, projecting a hypothetical, approximate membrane plane onto the structure (using the partially resolved TM helices) would indicate extensive overlap between the membrane and the bound Fab domain(s) (Fig. 1E). Constructing a membrane around the Fab-bound HA, involving both transmembrane and multiple, large peripherally interacting protein parts poses a challenge to conventional modeling techniques. To address this issue, we designed a novel strategy involving additional steps to allow the embedding membrane morph optimally into the experimental structure for the complex. The final model and its simulations allowed us to capture direct, functional interactions between the Fab and the surrounding lipids.

**Figure 1:**
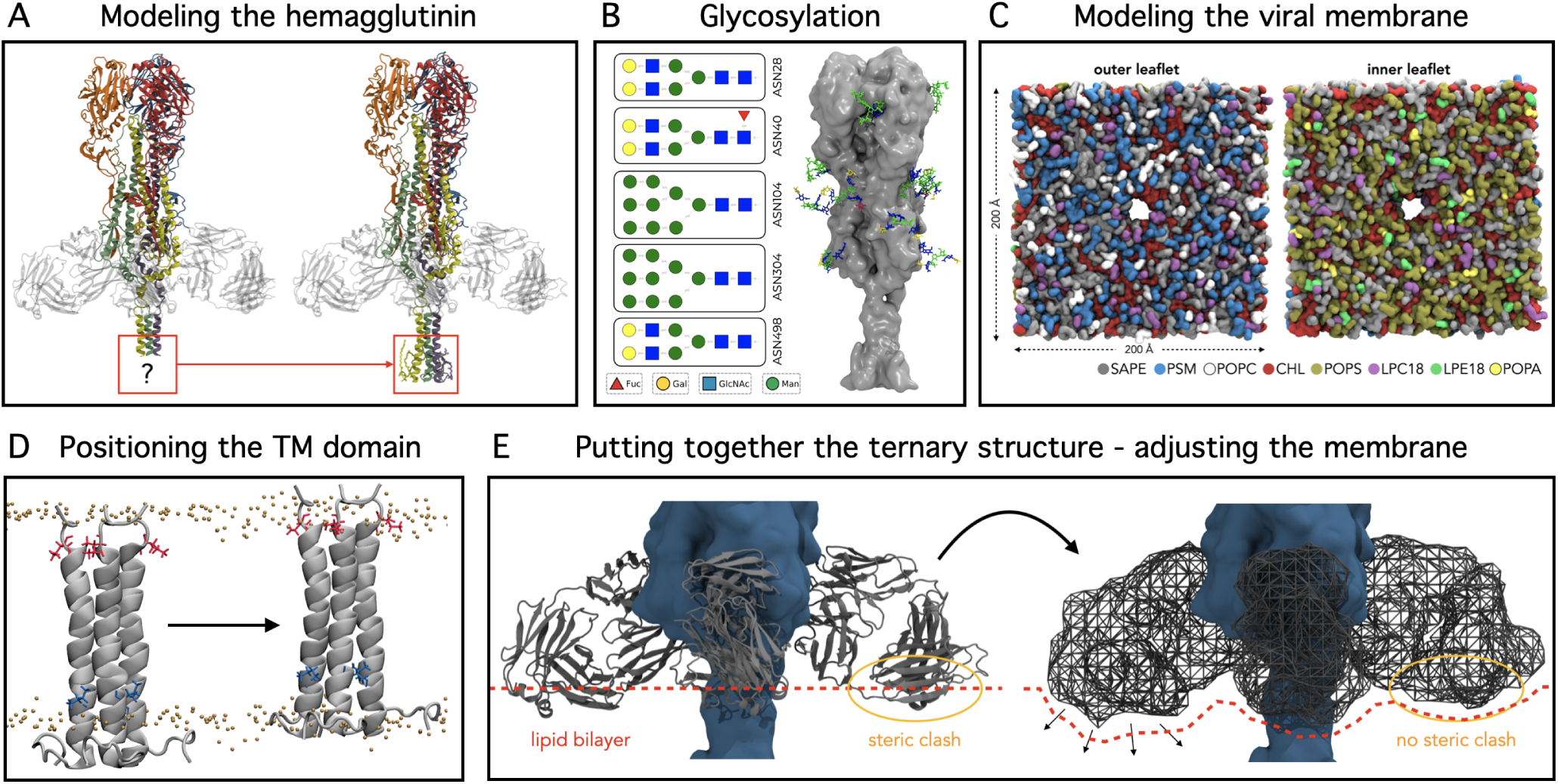
Overview of the protocol used to model influenza hemagglutinin (HA)-membrane and Fab-membrane interactions. (A) Comparison of the starting cryo-EM structure (left) and the constructed complete model (right) of HA wherein the missing transmembrane (TM) domain and cytoplasmic tail were modeled and palmitoylated. (B) Types of N-linked glycans modeled in the full HA structure. (C) A biologically accurate and asymmetric viral membrane which will be used to embed the Fab-bound HA. (D) Before putting the pieces of the ternary HA/Fab/membrane structure together, the depth of the modeled TM domain in the membrane was optimized via independent equilibrium simulations. (E) Given the highly curved micelle used in the cryo-EM experiment, placing the resulting Fab-bound HA into a planar viral membrane results in significant steric clashes with lipids. Therefore, the antibodies were initially represented as a repulsive grid potential (without an explicit representation), in order to allow neighboring lipids to freely diffuse and adopt a clash-free configuration.

HA is composed of a membrane-distal, receptor-binding domain (HA1) and a highly conserved, membrane-proximal, stem region (HA2)^13,27^. The membrane-embedded part of HA2 (the TM domain), and the following palmitoylated cytoplasmic tail (CT) play important roles in the overall conformations and fusogenicity of the HA trimer, as well as in the immune response to it^28^. Due to experimental limitations, even the best available cryo-EM structure (PDB ID: 6HJQ^17^) contains only a partially resolved TM domain, missing completely the CT (Fig. 1A). To complete the structure, we first utilized the transform-restrained Rosetta protein structure prediction server^29,30^ to construct the TM and endodomain. We then added palmitoyl groups at the cysteine-rich clusters of the CT region, which is one of the factors determining complete TM embedding in the membrane^31^.

The surface of the influenza HA is known to be heavily glycosylated^6^, and the glycans mediate various processes such as viral attachment to and entry into the host cell^32^. Recent mass spectrometry studies have thoroughly identified the glycan sites as well as their compositions for an HA variant from an H1 strain that is highly homologous to the HA studied here^33^. We thus modeled the complete glycans based on the mapping to the HA of the A/California/04/2009 (H1N1) viral strain determined by Thompson *et al.*^33^ while preserving the atomic coordinates of partial glycans already resolved in PDB:6HJQ (Fig. 1B). Throughout the subsequent MD simulations, the glycans were free to move and fully interact with their surroundings, thus contributing to the overall conformational dynamics of HA (Fig. S1).

### Adapting an explicit virion membrane for placement of Fab domains

The lipid composition plays a crucial role in the Fab’s interactions with the membrane. To best mimic the lipid profile of an influenza viral membrane, we adopted the lipid composition of H1N1 virion quantitatively measured with mass spectrometry and fluorometry (more details in Methods)^34,35^ when constructing the bilayer. Thus, we built an asymmetrical viral membrane patch including diacylphospholipids, lysophospholipids, sphingolipids, and cholesterol (Fig. 1C). The composition of the asymmetric bilayer was designed following these considerations: a) the molar ratio of phospholipids was adopted from the lipidomic study on the A/PR/8/34 (H1N1) virus^35^ given its close resemblance to the strain investigated here, with the percentage compositions scaled down to ~55% to allow for addition of cholesterol; b) both leaflets contained ~45% cholesterol, assuming its rapid cross-bilayer equilibrium due to flip-flop motions^36^; c) selective placements of anionic lipids only in the inner leaflet, and sphingolipids and phosphatidylcholines only in the outer leaflet, assuming such asymmetry is maintained in nascent viral particles, which inherit their lipid composition from the plasma membrane of the host cells; and, d) each type of head group is paired with only one tail configuration of the most common chain length and unsaturation characterized in the lipidomic study^35^. Accordingly, the bilayer was composed of SAPE:PSM:POPC:lysoPC:CHL (24:18:8:6:44) and SAPE:POPS:POPA:lysoPC:lysoPE:CHL (23:25:2:3:3:44) in the extracellular and cytoplasmic leaflets, respectively.

The position of the TM domain in the membrane is a key determinant of the extent and mode of Fab-lipid interactions. Therefore, before integrating the full model of Fab-bound HA, we placed a truncated TM/CT domain model in the bilayer and performed equilibrium simulations to obtain its optimal placement in the membrane (Fig. 1D).

The TM-containing cryo-EM structure was solved in a micelle environment. Placement of a planar bilayer patch over the structure resulted in substantial steric clashes between the HA-bound Fab domains and the bilayer lipids, which could not be resolved with simple energy minimization or even with regular MD simulations. We therefore devised a protocol to model the peripherally interacting antibodies together with the membrane, such that the lipids had maximal freedom to adjust to and accommodate the experimental structure of the complex.

First, to represent the resolved antibody structure, we replaced the Fab domains by spatial repulsive potentials (more details in Methods) (Figs. 1E, and S2A)^37^ in which lipids could easily move and adjust their positions without entanglement to atoms originating from the explicit presence of the antibodies. We then performed simulations with these potentials until the membrane adjusted fully to the experimental structure. Additionally, we adapted a non-periodic bilayer that would allow more readily for adjustments in both global and local lipid curvatures around the Fab insertion area (Fig. S2B-C) as well as some lipid flipping between the two leaflets.

The modeling and simulations were repeated for systems with varying numbers of Fabs since the minimal required stoichiometry for FISW84 to neutralize HA is unknown. We randomized lipid arrangements for each system to minimize biasing and performed 8 independent simulations each for 1 *µ*s.

### Antibody residues responsible for membrane binding

To characterize the specific contacts between the antibody and the membrane lipids, protein-lipid interactions were counted using all heavy atoms within 3.5 Å. We analyzed 12 bound Fabs among 6 different simulation runs, 2 replicates for each singly, doubly, or triply Fab-bound HA. Notably, none of the antibodies dissociated from the membrane in any of the simulations. On the contrary, Fab residues spontaneously anchored deeper into the membrane during the simulations stabilizing membrane binding. Aggregating all contacts across all simulations we identified the residues contacting the membrane most frequently (Fig.4A). Consistent with the expectations from the bound HA orientation, the heavy chain (orange in Fig.4B) formed the most extensive interface with the membrane. Additionally, long side chains in the light chain (purple in Fig. 4B) also demonstrated membrane anchoring.

To better describe the interactions between the Fabs and the lipids, we calculated their interaction energies for each Fab focusing on the previously identified membrane-binding residues as highlighted in Fig. 3A. The analysis showed that residues K126, R19, and K209 exhibited particularly strong interactions with the membrane (Fig. 3B), primarily due to electrostatic interactions between these charged residues and lipid head groups^38^.

We corroborated the findings from the simulations by assessing the neutralization activity of various designed FISW84 mutations against the HA stem of influenza A/PR/8/34 (H1N1) virus, which is similar in sequence to influenza A/duck/Alberta/35/76 (H1N1) virus (Fig. 3C). First, we aimed at modifying the electrostatic properties of the interfacial residues by applying charge-reversal mutations (e.g., R19E and K126E in the heavy chain, and K127E in the light chain). Charge-reversal mutations such as R19E increased the half-maximal effective concentration (EC_50_) by fourfold, suggesting that electrostatic interactions between these residues and membrane are important for stable binding of the antibody to HA (Fig. 3D). Next, we mutated polar residues to alanine (e.g., S124A and T125A in the heavy chain), to determine the role of hydrogen bonding. Hydrogen bonds between polar residues and the membrane turned out to be important for stable binding and neutralizing activity of antibodies, as exemplified by the T125A mutation in the heavy chain. Then, to potentially improve the membrane binding of the antibody, we mutated some of the interfacial polar residues to aliphatic or aromatic ones (e.g., S124L/W, T125L/W, N213W), in an attempt to create a deeper membrane anchor and enhance membrane insertion. However, the EC_50_ of these hydrophobic point mutations shows they do not improve viral neutralization (Fig. 3D), possibly due to reasons unrelated to their membrane interactions. It is likely that multiple point mutations are required to create a membrane-anchoring hot spot on an otherwise hydrophilic surface. Lastly, to investigate the impact of potential snorkeling effect of basic residues in the membrane, we mutated lysines and arginines to alanines (e.g., R19A, K126A). While salt bridges only take into account oppositely charged groups, snorkeling also includes the long hydrophobic chains of lysines and arginines that interact with hydrophobic core of the membrane. The increase in EC_50_ of the K127A mutation in the light chain suggests that the snorkeling effect of positively charged residues in the wildtype system contributes to stable antibody binding.

We noticed some of the lipids contacting the HA-bound Fabs are phosphatidylserines (PS), which is a negatively charged phospholipid predominantly found in the inner leaflet of the plasma membrane and was not present in the upper leaflet of the initial model. The membrane in our simulation system resembles a bicelle where the two leaflets are connected; thus, PS from the lower leaflet may diffuse to the upper leaflet and become Fab-bound. Since the viral envelope forms by budding from the plasma membrane, PS is expected to be in the inner leaflet. However, they can still be discovered in the outer leaflet upon viral infection for the following reasons. First, a viral-infected cell often activates lipid scramblases^39^, which leads to PS exposure at the cell’s surface^40^. Secondly, for the viruses which survive longer periods of time in the extracellular environment, due to the absence of ATP/flippase in the virions, the viral membrane would slowly lose its leaflet asymmetry by simple diffusion. While the precise exchange rate or concentration of PS lipids remain uncertain, in addition to our *in vitro* experiments, our computational models involving lipid flipping through diffusion have captured this phenomenon (Fig. S4A). Throughout our simulations, basic residues such as K127, K209, and R213 closely interact with the negatively charged PS headgroups (Fig. S4B). Although our design did not include PS in the outer leaflet, their interaction with antibodies can be linked to cross-leaflet migration due to the bicelle-like bilayer structure in the simulations.

### Structural insights from HA-antibody-membrane simulations

Our simulations revealed notable, functionally-relevant dynamics in HA, specifically in the flexible linkers connecting the TM domain to the ectodomain, in the receptor-binding head domains (HA1), and, most importantly a significant breathing motion in the head domains.

Due to the flexibility of the linkers connecting the TM and ectodomain, HA exhibited a substantial tilting motion (see Methods for the definition). Our simulations revealed development of tilt angles, ranging ~20-50*^◦^* (Fig. 2A). The binding of Fabs significantly hindered the HA tilting motion, particularly when more than one Fab was bound to HA (Fig. 2A). These findings are in close agreement with different TM orientations in the two cryo-EM structures of HA, where Fab-free HAs exhibit a tilt of 52*^◦^* ^17^, whereas Fab-bound HAs display only a 20*^◦^* tilt. Our findings regarding Fab-free tilt angles are also in line with HA tilts reported for simulations done in the context of the whole-virion^41^.

**Figure 2:**
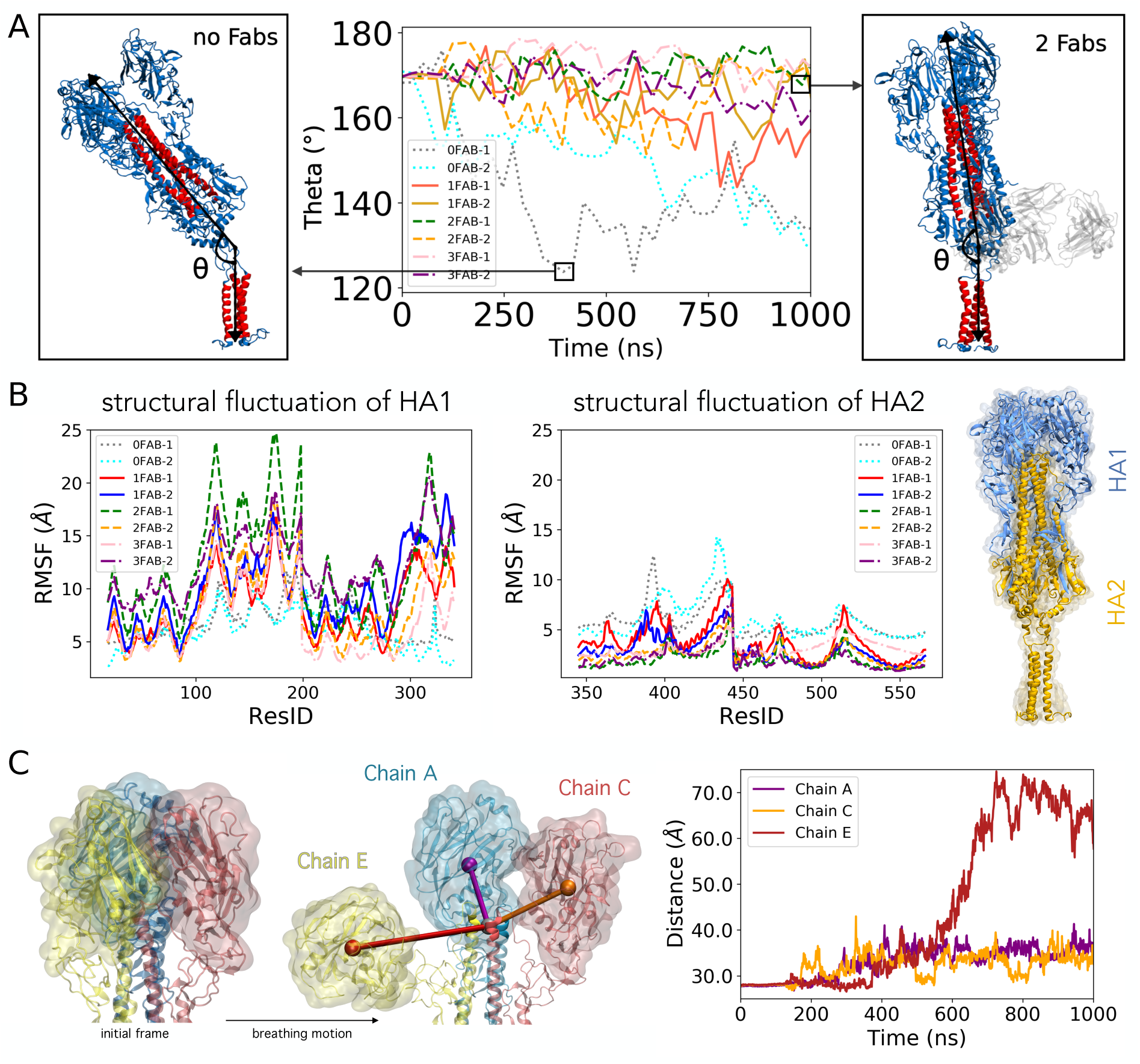
Antibodies govern the structural dynamics of HA. A) Tilting of the ectodomain is prevented by the stem-binding antibody. The TM domain exhibits a range of tilt motions deviating it from its threefold symmetry axis. The tilt is diminished in the Fab-bound HAs (the naming convention, e.g., for system 1 with 3 Fabs is 3FAB-1). B) Comparing the RMSF of the HA1 and HA2 domains of HA in the presence of different numbers of Fab domains for each system. For HA1, the systems without Fab are much more mobile than the Fab-bound ones. In contrast, for HA2, Fab-bound systems are more stable. C) Activation motion of the head domains in HA. The left panel presents an illustration of the HA head domains transitioning from their initial ‘closed’ state to an ‘open’ configuration. In the right panel, we provide a representative time series demonstrating the change in the distances between the center of mass (COM) of each head domain (purple, orange, and yellow spheres, respectively) and the COM of the tip of the long alpha helices in the core of the structure (gray sphere).

As far as HA1 and HA2, we observed significant conformational changes within the head domain, while the stalk domain remained stable during the simulations. Root-mean-square fluctuation (RMSF) analysis of HA1 and HA2 (Fig.2B) confirms that HA1 exhibits greater mobility compared to HA2. The HA1 domain, which is responsible for the glycoprotein’s breathing motion, showed reversible separation (shown in Fig. S5 bottom left panel) in its three globular domains (Fig.2C; see section below for detailed discussion on this motion).

Interestingly, comparing Fab-bound systems to the Fab-free control systems, we observe that in the presence of Fabs, HA1 underwent more pronounced conformational changes, while HA2 experienced more diminished structural fluctuation (Fig. 2A). This might be one of the factors contributing to the role of FISW84 in neutralizing HA by stabilizing the HA2 domain, which is critical in HA’s fusion-related conformational changes^9^.

The Fab elbow angle, serves as a valuable descriptor for the overall topology of the Fab fragment^42^, providing insight into the relative disposition of the variable and constant domains. It is frequently measured in Fab structures as a metric for Fab flexibility^43^, and studies have demonstrated the importance of enhancing its flexibility in improved recognition by diverse antigens^44^. Following Ferná ndez-Quintero *et al*.^44^, the elbow angle is defined between the pseudo-two-fold axes that relate the two pairs of domains (VH, VL, CH, CL), as illustrated in Fig. S6A. Throughout the simulations, the elbow angles were found in agreement with X-ray structures studied in Ferná ndez-Quintero *et al*.^44^ for hundreds of other Fab domains. This agreement supports the idea that the Fabs in our simulations maintained their structural integrity and were not significantly distorted by the membrane (see Fig. S6B). Furthermore, comparing systems with 1-3 Fabs, we found no significant differences in the elbow angle (Fig. S6C).

The Fab domains in all simulated systems exhibited a gradual further insertion into the lipid bilayer while maintaining a consistent structure and interaction with HA. When atomistic representations of Fabs were introduced back into the initial volumetric grid, the membrane was observed to progressively approach the Fabs. Initially, there was little to no contact or only minimal interaction between the membrane and the Fabs, but over time, the Fabs embedded more and more in the membrane, as depicted in Fig. S7. This embedding occurred rapidly (within a few nanoseconds), and persisted throughout the entire simulation, as supported by their maintained interaction (Fig. S7).

### Opening of HA head reveals cryptic epitopes

Recent discoveries have revealed a concealed and consistent section within the HA head of H1 (A/Solomon Islands/3/2006), holding promise for the development of a universal flu vaccine^45^. This concealed epitope is the target of trimer interface antibodies^45–47^, which provides broad protection against a number of flu strains. However, the epitope remains hidden when HA is in its closed form (e.g., in the cryo-EM structure) and its exposure relies on HA’s structural “breathing” motion.

The exposure of this epitope for trimer interface antibodies was reported by Casalino *et al.*^41^ in a simulation study performed for a crowded virion environment. In the span of 500 ns, their simulation indicated an asymmetrical motion within the head domains, with two of the the globular components displaying pronounced splitting while the third remaining closed.

Aligned with those findings, our *µ*s simulations demonstrate that the epitope for trimer interface antibodies indeed becomes reachable as the HA head domain undergoes a significant breathing motion (Fig. S5). In order to assess the accessibility of the hidden site, we superimposed HA in our systems with the crystal structure of the head domain of H1 bound to a representative trimer interface antibody, FluA-20 Fab (PDB ID: 6OC3^45^), to show the accessibility of this hidden site as the head domain opens up (Fig. 5, see details in Methods). Remarkably, irrespective of the number of Fabs bound to HA, the majority of instances across half of our simulations reveal HA conformations to which FluA-20 can bind. Notably, the relationship between the head-splitting distance and FluA-20 successful docking is not necessarily linear, suggesting that a twisting motion of the HA head domains through the flexible linking loops also contributes to rendering the hidden epitope accessible.

## CONCLUDING REMARKS

Here, we report the first membrane-bound model for HA in complex with one of its stem-binding Fab. We model and simulate HA-Fab-membrane complexes with varying numbers of Fab domains and analyze the direct interactions between the Fab and the lipids of the membrane to investigate the importance of the membrane interaction in neutralization activity of the antibody. The importance of specific interactions at the membrane interface is validated by experimental mutagenesis and virus neutralization assays. The reported simulations capture large-scale conformational changes of the head domain and a stable stem. Depending on the number of Fab domains bound to HA, significant tilting of HA is also observed, in agreement with the cryo-EM density maps for the protein. The variable domain of the Fabs is found to be highly stable compared to the constant domain, indicating a healthy antibody-antigen interaction. Additionally, the simulations capture dynamic and transient accessibility of the cryptic epitope for trimer interface antibodies during HA head breathing motion.

## MATERIALS AND METHODS

### Construction of full-length HA

As a first, necessary step, full-length HA was constructed (as shown in Fig. 6) based on a recent cryo-EM structure of influenza HA (strain A/duck/Alberta/35/1976 H1N1) in complex with FISW84 Fab fragments (PDB ID 6HJQ^17^). The missing flexible loops were modeled and the altered residues in the PDB entry were reverted back to wildtype (V125I, K236Q, L547W) using PDBfixer (an OpenMM tool)^48^, referencing the protein sequence from Uniprot ID P26562 (HEMA I76A4) where we follow for our residue numbering: HA1 (M1–R344) and HA2 (G345– I566) in a continuous sequence numbering. The omitted loop in the HA-bound FISW structure, which corresponds to S137–G142 in the reference IGG sequence (Uniprot entry P01857), was not modeled. Note that the loop is present in the F_ab_ constructs used in the neutralization experiments described later.

A total of 18 disulfide bonds were introduced between cysteine pairs. Following the Uniprot ID P26562 numbering, these cysteine pairs were C59–C292, C72–C84, C107–C153, and C296– C320 in HA1, C488–C492 in HA2, C21–C481 between HA1 and HA2, as well as 12 pairs within the Fab fragments (C22–C96 and C143–C199 in each heavy chain, and C23–88 and C135– C195 in each light chain). The helical TM domain of the cryo-EM structure, defined as residues Q529–M554^17^ is only resolved up to residue G548. Without a suitable template, the rest of the helix followed by the loop was modeled using the transform-restrained Rosetta protein structure prediction server^29,49^, including the CT, and residues C555–I566. Residues Q529–M554 were used to align the TM homology model with the crystal structure for each monomer, completing the model. Then, in the endodomain, 3 palmitoyl groups were adopted in each monomer (connected to C555, C562, and C565) in the cysteine-rich clusters of the C-terminal region using CHARMM-GUI^50,51^.

Partially resolved glycans were completed by extending them using the mapping of HA-glycosylations of A/California/04/2009 (H1N1) identified in a recent mass spectrometry study^33^. Five glycosylation sites in each HA monomer were modeled, resulting in 15 N-glycans (Fig. 1B) using the PSFGEN package from VMD (Visual Molecular Dynamics)^52^. Glycan topologies from the crystal structure and the model were examined before simulation, and any *cis* to *trans* conversion at the amide bond (C2-N2) of N-acetylglucosamine (GlcNAc)^53^ were detected and corrected using Conformational Analysis Tools^54^.

### Embedding into a virion membrane for juxtaplacement of Fab domains

For placement of the complete TM domain in a membrane, a 200 200 Å ^2^ membrane patch was chosen, with a lipid composition mimicking PR8 influenza viral membrane, which has been determined with mass spectrometry and fluorometry^34,35^. The bilayer was composed of 1-stearoyl-2-arachidonoyl-sn-glycero-3-phosphoethanolamine (SAPE), palmitoyl-sphingomyelin (PSM), 1-Palmitoyl-2-oleoyl-sn-glycero-3-phosphocholine (POPC), 1-palmitoyl-2-oleoylglycero-3-phospho-serine (POPS), palmitoyl-oleoyl-phosphatidic acid (POPA), 18:0-lyso-phosphatidylcholine (Lyso-PC), 18:2-lyso-phosphatidylethanolamine (LysoPE) and cholesterol (CHL) at molar ratios of SAPE:PSM:POPC:lysoPC:CHL at (24:18:8:6:44) and SAPE:POPS:POPA:lysoPC:lysoPE:CHL at (23:25:2:3:3:44) in the extracellular and cytoplasmic leaflets, respectively. The lipid bilayer was initially built using CHARMM-GUI^50,51^. However, due to the lack of the needed lipid topologies for 18:2-LysoPE and 18:0-LysoPC, these lipids were initially modeled as LPC14 (1-myristoyl-2-hydroxy-sn-glycero-3-phosphocholine; 14:0 LysoPC) and LPC16 (1-palmitoyl-2-hydroxy-sn-glycero-3-phosphocholine; 16:0 LysoPC) in CHARMM-GUI and later replaced with the corresponding lysolipids using PSFGEN. To determine the relative positioning of the TM domain in the membrane, equilibrium simulations were performed (150 ns) for the TM trimeric model first in isolation, where the depth of the TM with respect to the bilayer was determined (Fig. S8).

The PDB structure of antibody-bound HA is resolved in a micelle, and its placement in a flat membrane resulted in steric clashes between the Fab domains and the bilayer. In order to allow for free changes in the membrane shape when bound to the protein complex, a non-periodic bilayer (with free edges) was adopted in the simulation box to allow free deformation of the membrane from planarity, which might be needed to accommodate the HA-bound Fab domains. Next, to avoid any lipid trapping between HA and the Fab domains during the initial membrane placement, the Fab domains were represented by a lipid-repulsive, void volume (as a grid map exerting a repulsive potential to the lipids) matching their experimental geometries relative to HA. The system was then simulated using the molecular dynamics flexible fitting (*mdff*) protocol for 17 ns (Fig.1E)^37^.

Thus 4 different systems were assembled for protein-membrane complexes including zero to three Fab domains bound to the HA trimer. Each system was then simulated in two independent simulation replicas. To avoid any bias from the initial lipid configuration and to improve sampling, for each simulation replica, 20% of the lipids in the bilayer were shuffled using the Membrane Mixer plugin (MMP) in VMD^55^ before the MD simulations.

### MD and MDFF simulations

The membrane-embedded, full-length HA system without Fab was solvated and ionized with 0.15 M NaCl, resulting in an antibody-free simulation system with approximate dimensions of 222 222 227 Å^3^ with 1.2 million atoms. During all later steps of modeling and equilibration of this system, with or without antibody, to maintain the experimentally resolved structures, all protein backbone atoms were restrained using harmonic potentials (*k* = 5 kcal/mol/Å^2^). The Fab-free system was equilibrated using a single 15-ns NPT simulation. The Fab-containing systems were prepared for production runs in 3 steps. First, after 2,000 steps of steepest descent minimization, a 15-ns, restrained equilibration under NPT conditions was performed. In Step 2, grid maps representing the shapes of 1, 2, or 3 Fab fragments were introduced to the system to exert repulsive forces on lipids/solvent and “carve” the volume needed for all-atom antibodies to be added later. In this step, the systems were simulated each for 17 ns. Finally, in Step 3, the atomic structures of 1, 2, or 3 Fab fragments were introduced back into the void volumes, resulting in system sizes of around 1.5 million atoms and approximate dimensions of 250 250 270 Å^3^. The final constructs were then minimized again for 5,000 steps and equilibrated for 100–200 ns with protein backbone restraints as described above, to allow for further relaxation of lipids and solvent around the experimental protein structures.

After the equilibration phase, all the restraints were removed, and a production run was carried out for 1 *µ*s for all 8 constructs. Simulations were performed using GPU-resident NAMD^56,57^ employing the fully atomistic CHARMM36m^58^ force field for proteins and CHARMM36 force fields for the lipids and cystein palmitoylations^59^, N-linked glycosylations^60^, and ions. The TIP3P model was used for water^61^. The simulations were performed as an NPT ensemble with the temperature and pressure maintained at 310 K and 1 bar using Langevin thermostat and barostat, respectively^62,63^. The SHAKE algorithm was used to constrain all bonds with hydrogen atoms. For the calculation of van der Waals interactions, a pairlist distance of 13.5 Å, a switching distance of 10.0 Å, and a cutoff of 12.0 Å were used. The Particle mesh Ewald (PME) method^64^ under periodic boundary conditions was utilized for the calculation of electrostatic interactions and forces without truncation. The first equilibration phase was carried out using a timestep of 1 fs and the subsequent steps and the production runs with a timestep of 2 fs.

### Simulation analysis

The tilt angle of each HA monomer was quantified by measuring the angle (*θ*) between the TM domain principal axis and the ectodomain’s long *α*-helix axis (Fig. 2A). The vector representing the axis of the ectodomain’s *α*-helix connects the centers of mass (COM) of the residues at the top (residues 422-426) to those at the bottom (residues 460-467).

The separation of the head domains was quantified by measuring the distance between the COM of each HA head and the COM of the top residues (419-423) in the three long *α*-helices (Fig. 2C).

A representative simulation snapshot of a triply Fab-bound system where one of the head domains (Chain E) drastically separated is shown in Fig. 2C, and the full analysis is provided in Fig. S5.

Interaction of the Fabs and membrane lipids was investigated by measuring the contacts between them. A contact was defined for any Fab heavy atom within 3.5 Å of any lipid atom (Figs. 4 and S4). Contacts (collected every 1 ns) were categorized based on the residues in Fab. A cumulative histogram was then generated using the data from all 12 individual Fab domains from the 1 *µ*s simulations.

The time series of interaction energies between specific Fab residues and the lipid bilayer (calculated using NAMD ENERGY) for all systems are shown in Fig. S3, with their average values plotted in Fig. 3B.

**Figure 3:**
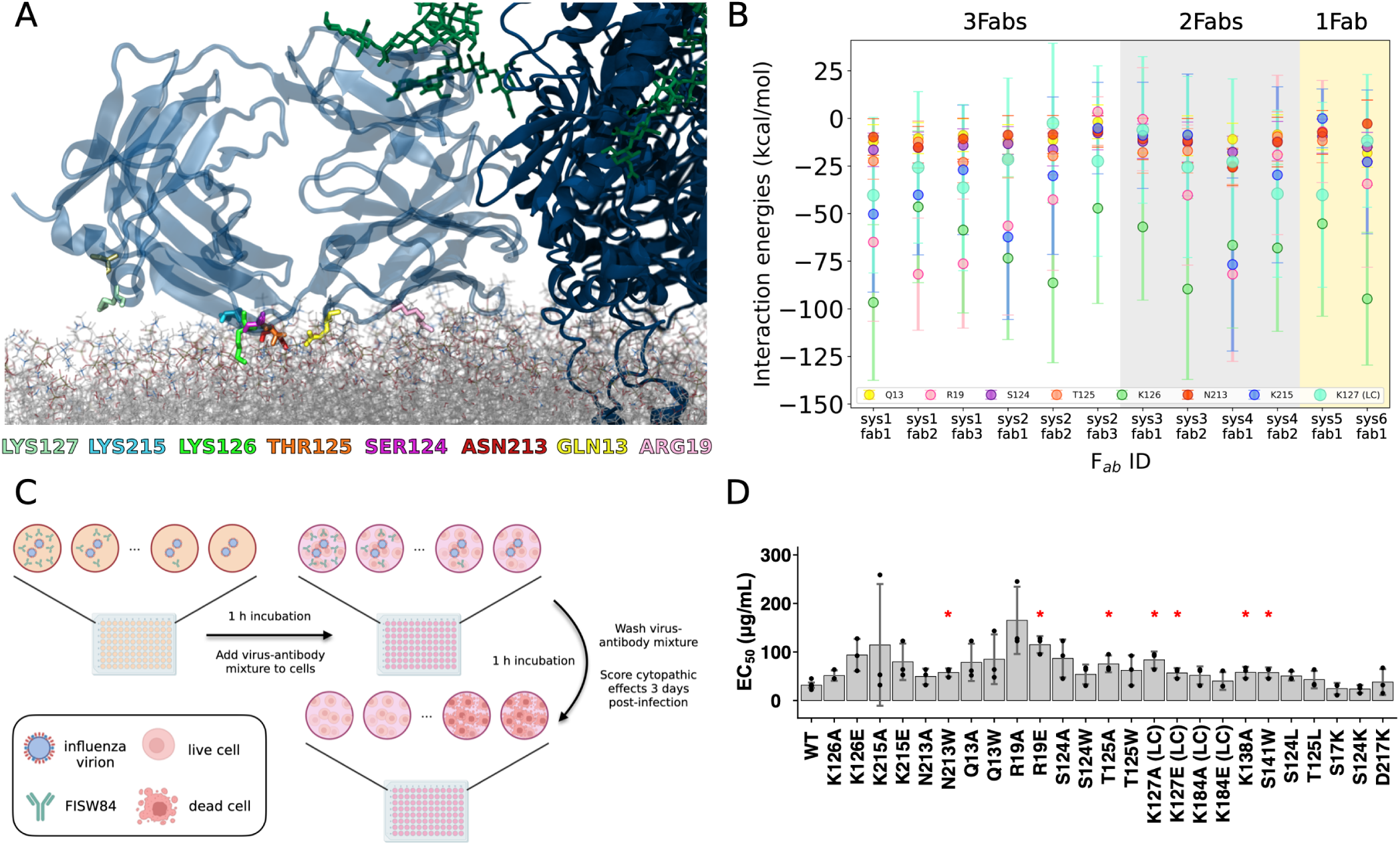
Characterizing membrane-binding residues. A) A snapshot of a membrane-bound Fab with highlighted interfacial residues interacting with lipid headgroups. B) Average interaction energies between select residues and membrane lipids in each simulation for systems with 1-3 Fabs. C) Summary of the virus neutralization assay. Two-fold dilutions of wild-type or mutant FISW84 antibody were incubated with influenza A/H1N1/PR8/34 virions for 1 h. The virus-antibody mixture was added to MDCK cells for 1 h, washed, and the half-maximal effective concentration (ED_50_) of wild-type or mutant FISW84 antibody was measured based on cytopathic effect two days post-infection. Figure created with Biorender. D) Mutagenesis experiments and virus neutralization assay show the importance of membrane-binding residues of influenza HA bnAb. Average half-maximal inhibitory concentration (EC_50_) values (bars) are measured for each FISW84 antibody mutant from three biological replicates. Each data point corresponds to the calculated EC_50_ from one biological replicate. Data represent mean standard deviation. Red asterisks indicate *p*-value ¡ 0.05 from a two-sided Student’s *t* -test. LC: light chain.

The accessibility of the hidden epitope of the head domain by the FluA-20 antibody is mediated by opening of HA^45^. To analyze this, we used a crystal structure of FluA-20 Fab complexed with the head domain of H1 (A/Solomon Islands/3/2006; PDB ID 6OC3) to assess whether the conformation captured within the simulations allowed for a clash-free docking of the Fab to the HA epitope. To accomplish this, the head domain from the PDB:6OC3 structure was superimposed onto each of the three individual head domains for every time frame throughout the trajectories across all eight systems. This procedure aimed to identify conformations in which the Fab could be docked to the epitope without giving rise to any steric clashes. The resulting “no clash” conformations are visualized as data points in Figure 5B.

**Figure 4:**
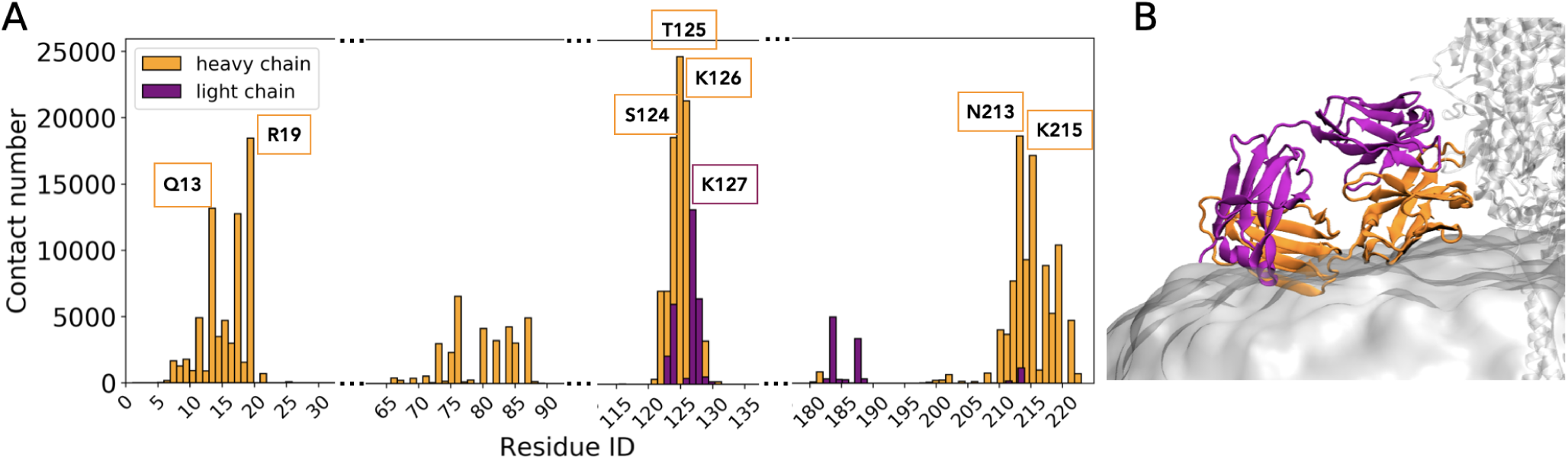
Membrane-binding residues of Fab. (A) The histogram displays the number of contacts (heavy atom pairs within 3.5 Å) between the Fab and the lipid bilayer. Residues that most frequently contribute to membrane binding are highlighted with orange (heavy chain) and purple (light chain) text boxes indicating their location. (B) A representative snapshot from a simulation of 3-Fab-bound HA highlighting the positioning of heavy (orange) and light (purple) chains with respect to the membrane (gray surface).

**Figure 5:**
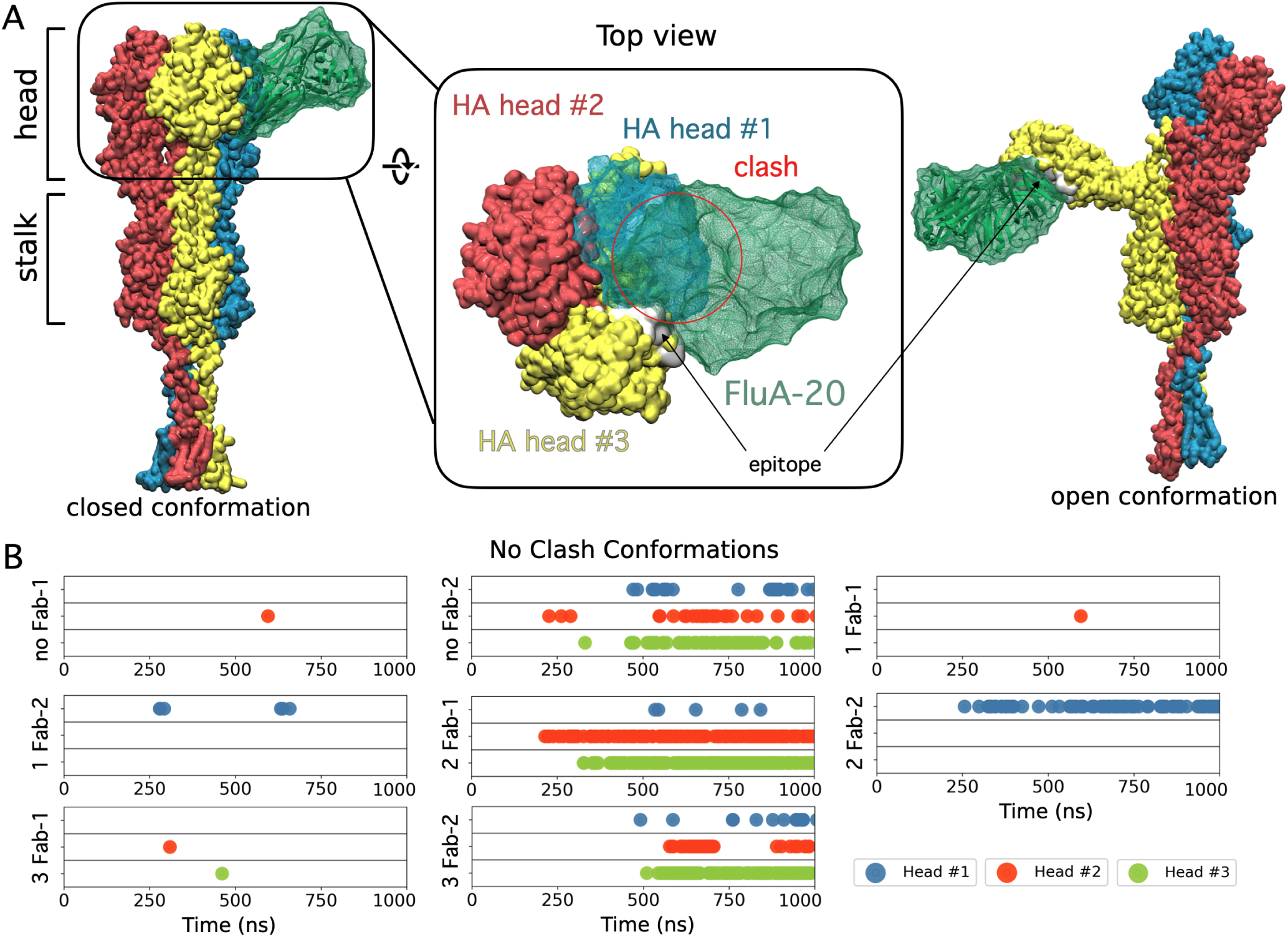
Exposure the hidden epitope through HA head domain splitting. (A) Docking of FluA-20 (green) to the cryptic epitope in the closed conformation (left/middle) of HA head domains (blue, red, yellow) results in significant steric clashes. Upon splitting of the HA heads and formation of the open conformation (right), the cryptic epitope becomes accessible and allows for binding of FluA-20 without any clashes. (B) Time series from all simulation systems depicting the formation of HA head conformations that allow for unobstructed binding of FluA-20 antibodies. These HA conformations, in which all atoms show a minimum distance of 1 Å to FluA-20 Fab), are shown as color points in the diagram.

**Figure 6:**
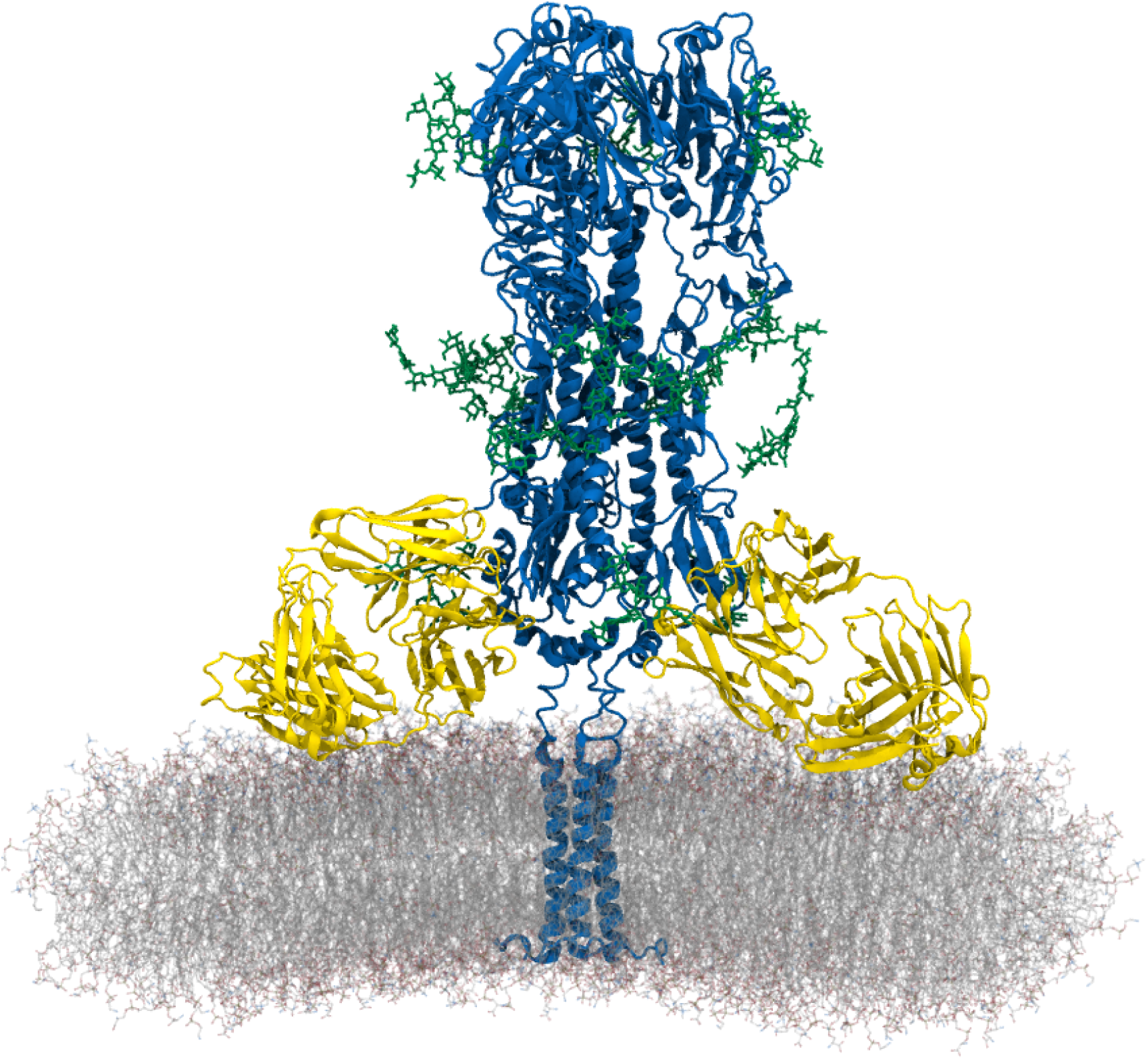
Ternary structure of antibody-bound, glycosylated HA embedded in a viral membrane. The modeled structure shows two FISW84 Fab fragments (yellow cartoon) bound to the HA (blue cartoon) epitope, which is embedded in a membrane patch representing the PR8 influenza viral membrane (lipids depicted as lines). N-glycans are displayed as green stick representations.

### Cell culture

Madin-Darby canine kidney (MDCK) cells (American Type Culture Collection, CCL-34) were grown and maintained in D10 medium – Dulbecco’s modified Eagle medium with high glucose (Gibco) supplemented with 10% v/v fetal bovine serum (Gibco), 1× non-essential amino acids (Gibco), 100 U/mL penicillin and 100 *µ*g/mL streptomycin (Lonza), and 1× GlutaMAX (Gibco) – at 37*^◦^*C, 5% CO_2_, and 95% humidity. Human embryonic kidney 293T (HEK293T) cells (American Type Culture Collection, CRL-3216) were grown and maintained in D10 medium under the same conditions. Expi293F cells (Gibco) were grown and maintained in Expi293F expression medium (Gibco) with shaking at 125 rpm, 37*^◦^*C, 5% CO_2_ and 95% humidity.

### Virus rescue and propagation

To generate a seed stock, eight pHW2000 plasmids encoding each of the segments of influenza A/Puerto Rico/8/1934 (H1N1) virus were transfected into a co-culture of HEK293T and MDCK cells (6:1 ratio) using Lipofectamine 2000 (Invitrogen) following the manufacturer’s protocol. Of note, influenza A/H1N1/Puerto Rico/8/1934 virus was used because of the highly conserved nature of the stem domain between influenza A/Puerto Rico/8/1934 (H1N1) and influenza A/duck/Alberta/35/76 (H1N1). Forty-eight hours post-transfection, media were aspirated and centrifuged at 300 × *g* for 5 minutes to remove debris. The clarified supernatant was used for propagation in MDCK cells to generate a working stock, which was then stored at 80*^◦^*C. The 50% tissue culture infectious dose (TCID_50_) of the inoculum was titered with MDCK cells.

### Plasmids

DNA sequences encoding the variable heavy chain and kappa light chain of FISW84 were codon-optimized for expression in human cells. Oligonucleotides encoding the variable regions of FISW84 heavy chain and kappa light chain were cloned into phCMV3 plasmids with a murine IGKV signal peptide in an IgG1 format. Sequences of these oligonucleotides are found in Supplementary Table 1. Mutagenesis was performed via PCR-based site-directed mutagenesis using either the heavy chain or kappa light chain as template, and forward and reverse primers listed in Supplementary Table 2. PrimeSTAR MAX DNA Polymerase kit (Takara) was used for PCR with the following settings: 98*^◦^*C for 10 s, 22 cycles of (98*^◦^*C for 10 s, 55*^◦^*C for 5 s, 72*^◦^*C for 30 s), 72*^◦^*C for 30 s. PCR products were then subjected to DpnI digestion (NEB) for 2 h at 37*^◦^*C. Subsequently, DpnI-digested products were transformed into chemically competent DH5*α Escherichia coli* cells. Plasmids were extracted with Miniprep kits (Qiagen), and sequences were verified via Sanger sequencing.

### Antibody expression and purification

Plasmids encoding the heavy chain and kappa light chain of WT or mutant FISW84 were transfected into Expi293F cells in a 1:1 molar ratio using an ExpiFectamine 293 transfection kit (Gibco) following the manufacturer’s instructions. Cell suspension was harvested 6 days post-transfection and the supernatant was recovered via centrifuging at 4000×*g* for 20 min at 4*^◦^*C. The supernatant was further clarified by filtration using a 0.22 *µ*m polyethersulfone membrane filter (Millipore). Subsequently, CH1-XL affinity beads (Thermo Scientific) were washed with MilliQ H_2_O and 1 × phosphate-buffered saline (PBS). Beads were then resuspended in 1×PBS. The clarified supernatant and washed beads were incubated at 4*^◦^*C overnight with gentle rocking. Then, flowthrough was collected, beads were washed with 1×PBS and incubated with 60 mM sodium acetate, pH 3.7 for 10 min at 4*^◦^*C. Antibodies were eluted and buffer-exchanged into 1× PBS using a centrifugal filter unit with a 30 kDa molecular weight cut-off (Millipore). Antibodies were sterile-filtered using 0.22 *µ*m cellulose acetate filters (Corning) and then stored at 4*^◦^*C.

### Virus neutralization assay

MDCK cells were seeded on 96-well plates in D10 medium at 8 × 10^4^ cells per well density, and incubated overnight at 37*^◦^*C, 5% CO_2_ and 95% humidity to reach ~100% confluency for infection the next day. Then, 100 TCID_50_ of influenza A H1N1/PR/8/1934 were mixed with two-fold serially diluted antibodies, with the highest and lowest concentrations at 500 *µ*g/mL and 0.98 *µ*g/mL, respectively, in 100 *µ*L of infection medium – minimum essential medium supplemented with 25 mM HEPES (Corning), 1× GlutaMAX (Gibco) and 1 *µ*g/mL tosyl phenylalanyl chloromethyl ketone (TPCK)-trypsin – at 37*^◦^*C, 5% CO_2_ and 95% humidity for 1 h. MDCK cells were washed with 1× PBS once and infected with 100 *µ*L of virus-antibody mixture for 1 h at 37*^◦^*C, 5% CO_2_ and 95% humidity. Then, inocula were discarded and replaced with 100 *µ*L of infection medium. Each antibody was assayed for virus neutralization with technical triplicates. Cytopathic effect as observed via cell death was scored 72 h post-infection. Figure 3D shows the schematic of the virus neutralization assay. Half maximal effective concentration (EC_50_) values were calculated in Prism v9 (GraphPad). Two-sided Welch’s *t* -tests were performed using R v4.1.1.

## RESOURCE AVAILABILITY

### Lead contact

Requests for further information and resources should be directed to and will be fulfilled by the lead contact, Emad Tajkhorshid.

### Materials availability

All plasmids generated in this study are available from the lead contact without restriction.

### Data and code availability

- Simulation trajectories have been deposited at Zenodo under the DOI 10.5281/zenodo.14014675 and is publicly available as of the date of publication.
- Any additional information required to reanalyze the data reported in this paper is available from the lead contact upon request.

## ACKNOWLEDGMENTS

This study was supported by the National Institutes of Health through the grants P41-GM104601, R24-GM145965, R01-GM123455, R01-AI167910, and DP2-AT011966. Simulations were performed using allocations on National Science Foundation Supercomputing Centers (ACCESS grant number MCA06N060), and Delta advanced computing and data resource which is supported by the National Science Foundation (award OAC 2005572) and the State of Illinois.

D.G.O. acknowledges support from the Beckman Foundation.

## AUTHOR CONTRIBUTIONS

D.G.O., T.J.C.T., N.C.W., and E.T. designed research; D.G.O., T.J.C.T., Q.W.T. and H.L. performed research; D.G.O., T.J.C.T., P.C.W., Z.G., and M.F. analyzed data; all authors wrote the manuscript.

## SUPPLEMENTAL INFORMATION INDEX

Figures S1-S8, Tables S1 & S2, and their legends in a PDF

