## Supplementary material for "Probing the Role of Membrane in Neutralizing Activity of Antibodies Against Influenza Virus": SI

### Supporting Information



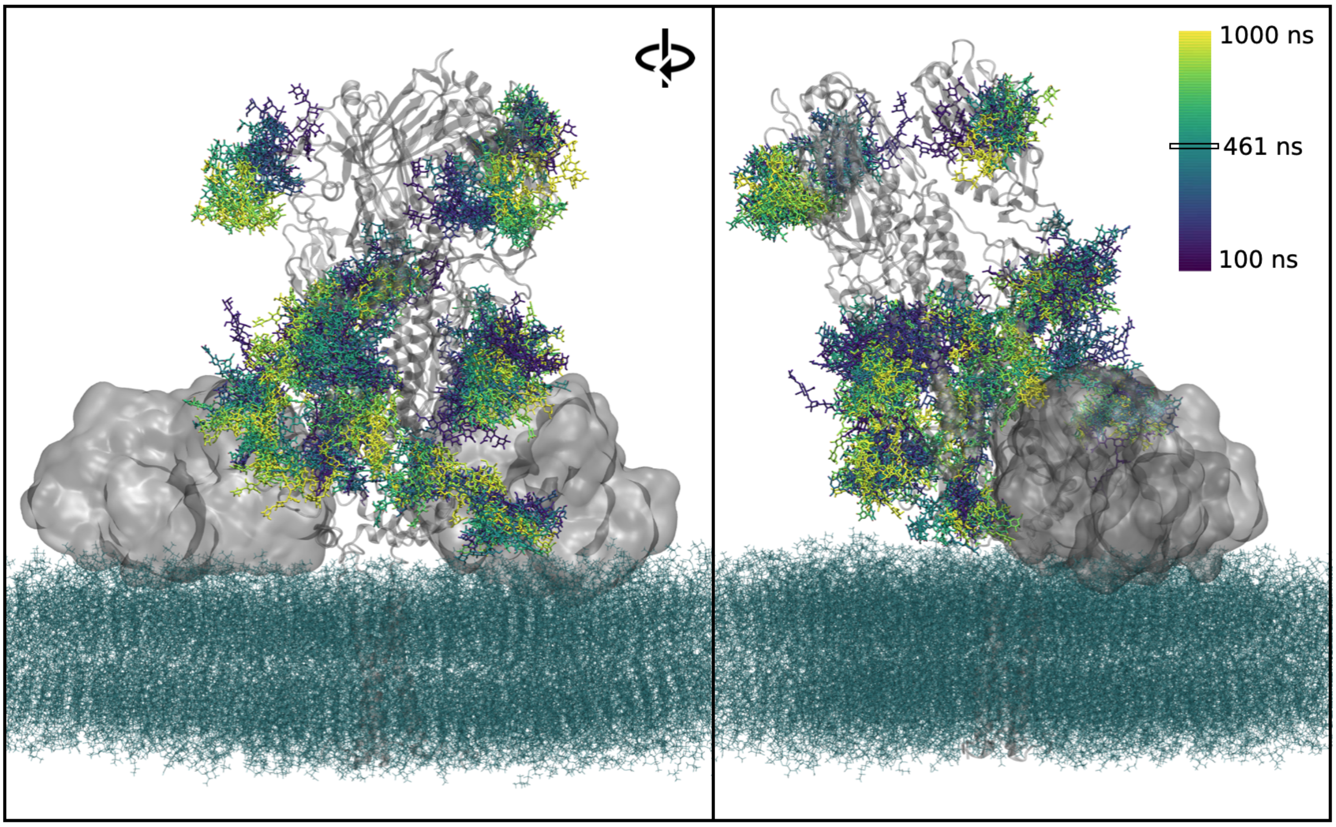


Figure S1: **Fluctuations of the glycans in HA.** The figure illustrates the motion of HA glycans (shown in licorice) over time, with each step depicted by a distinct color, according to the color bar. The displayed protein snapshot corresponds to frame at 461 ns, where Fabs are shown as transparent surface representation, the membrane as cyan lines and HA as transparent cartoon representation. The snapshot sourced from the system features doubly-bound Fabs captured from two different angles of view.

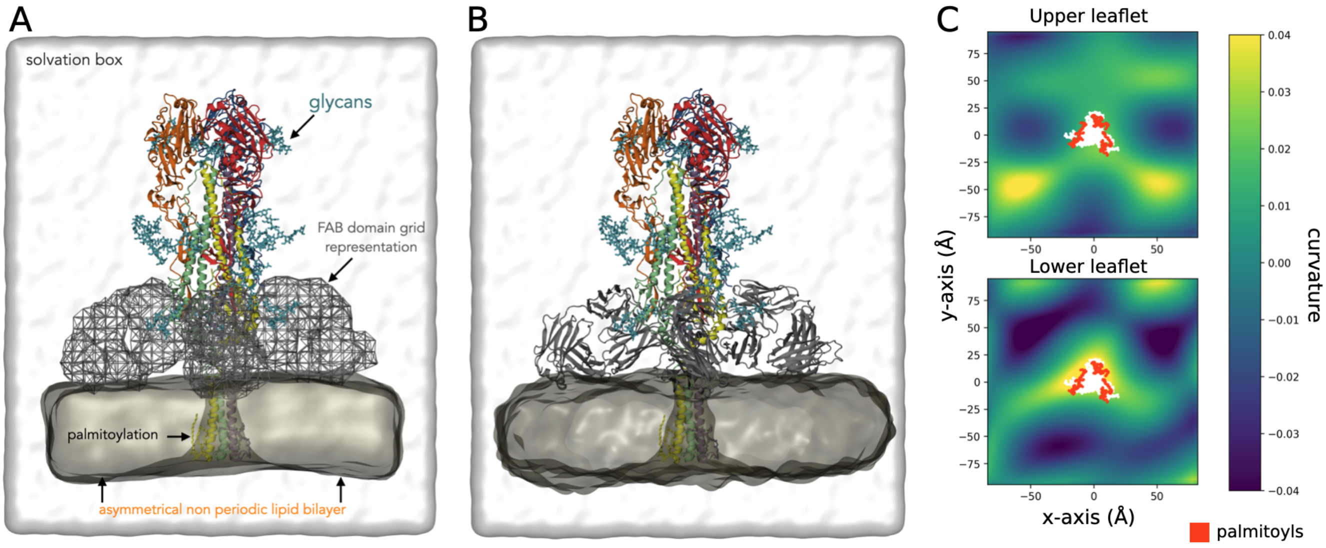


Figure S2: **Overall representation of the initial setup.** A) Initial simulation setup with a nonperiodic bilayer and Fabs represented as grid meshes exerting repulsive potential. B) Representation of the first snapshot from the equilibrium simulations after Fab atomistic placement and C) mean curvature of the membrane in B in the outer and inner leaflets. Curvature profiles in the center are affected by 3 Fab around palmitoylated (red) TM domains (white).

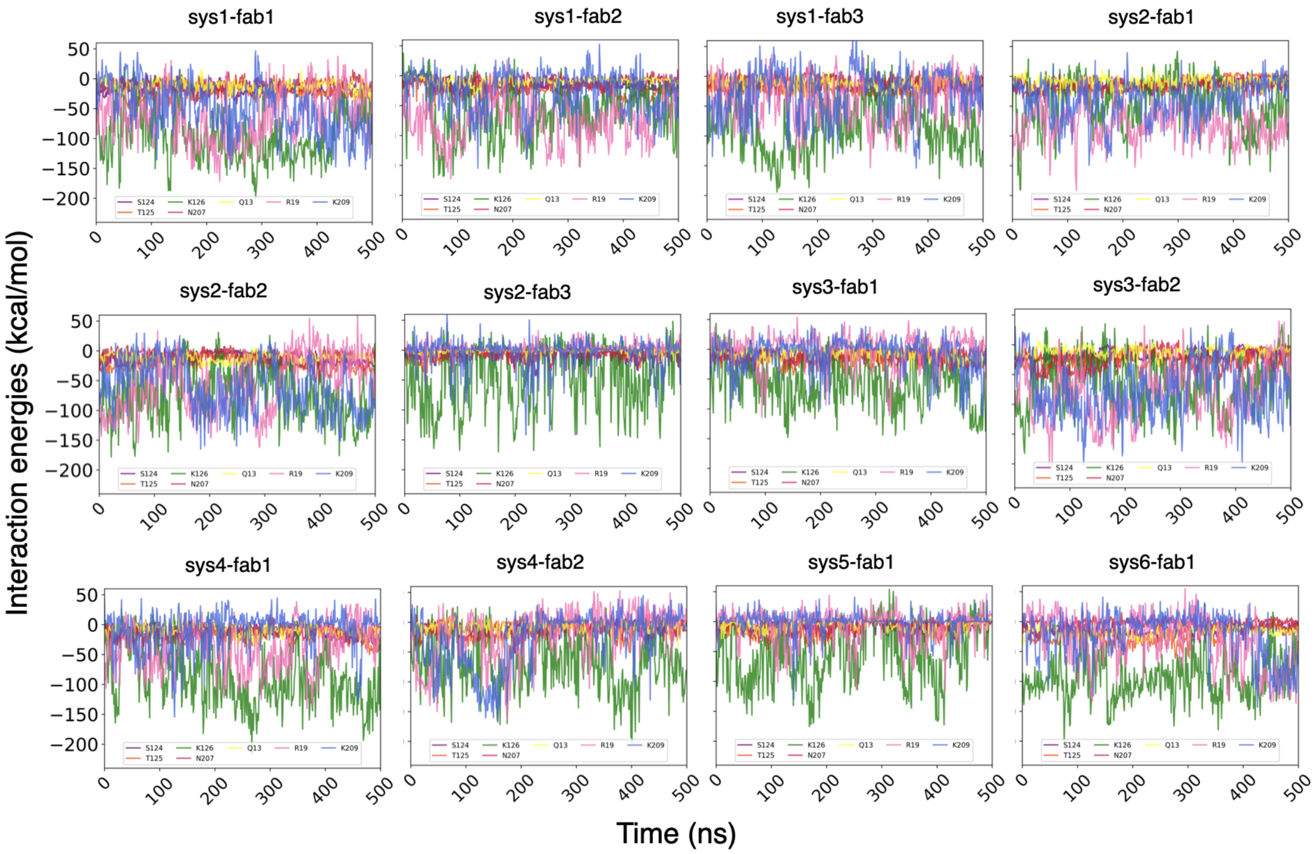


Figure S3: **Fab-membrane interaction energy per selected replica.** The time series for the interaction energy, as the sum of nonbonded interactions between select interfacial residues in the membrane-binding interface of Fab and the lipid bilayer are shown for all the systems individually.

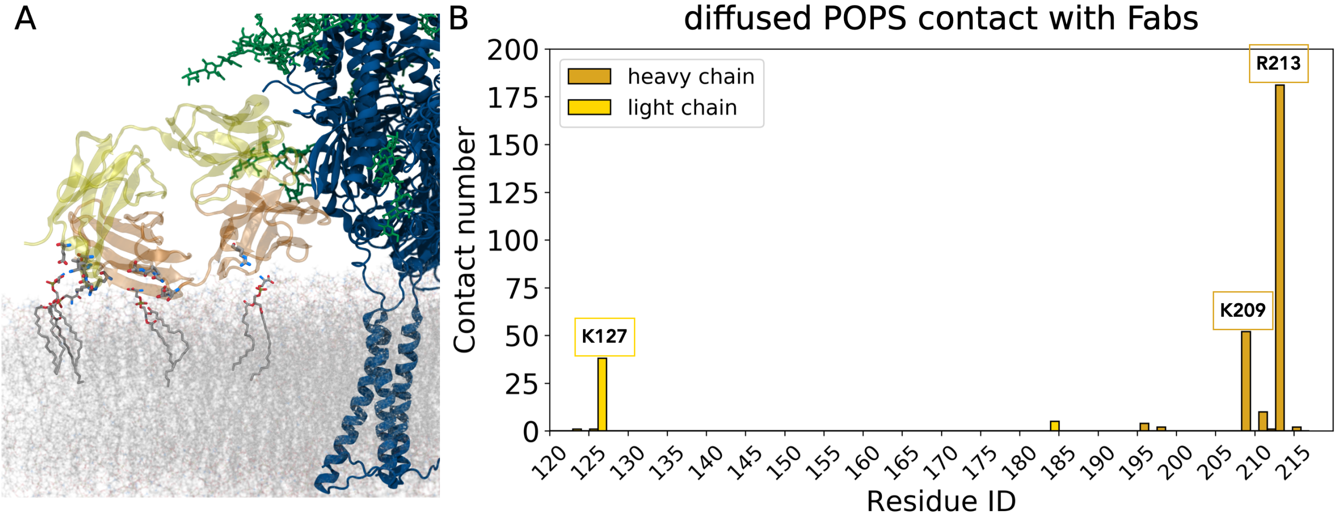


Figure S4: **Membrane POPS interacts with Fabs.** A) A snapshot showing the Fabs (light chain in transparent yellow, heavy chain in transparent orange) residues interacting with the POPS lipids (licorice representation). B) The contacts between the Fabs and the POPS lipids were measured throughout the simulations (see calculation details in Methods). A cumulative histogram from all 12 Fab domains is plotted using the contacts from the 1 $\mu$s simulations.

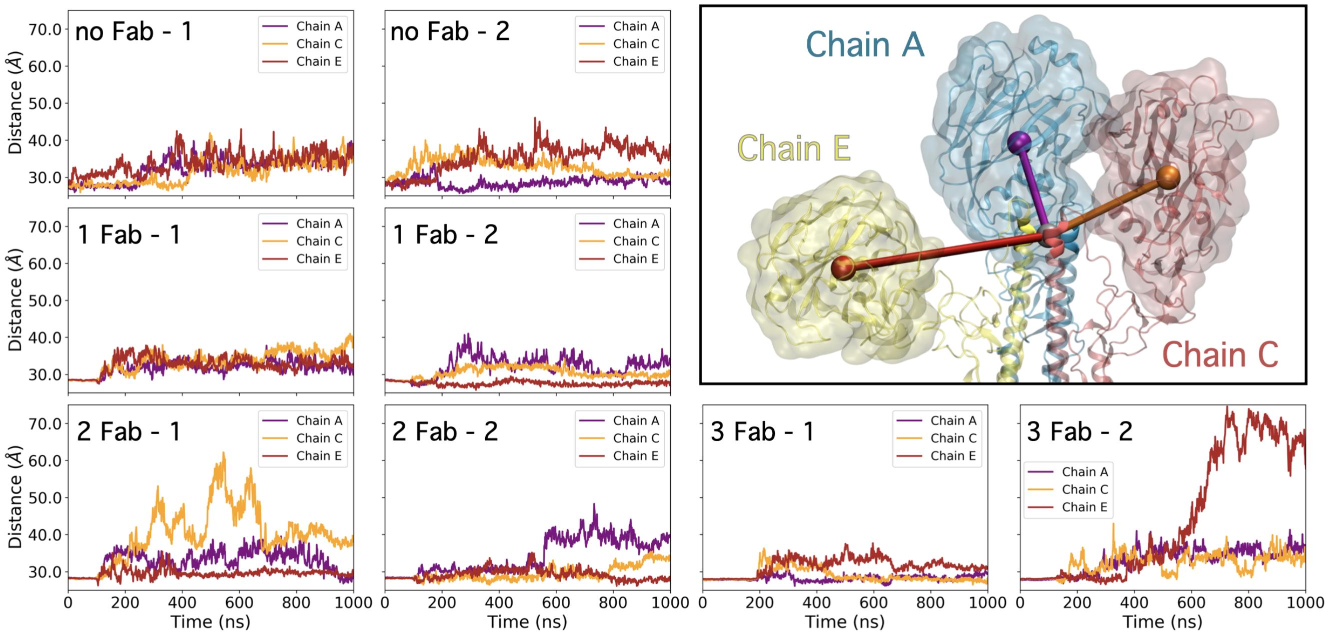


Figure S5: **Breathing motion of the HA head domains per trajectory.** The plot depicts representative time series of the distances between the center of mass (COM) of each head domain (purple, orange, and yellow spheres, respectively) and the COM of the tip of the long $\alpha$ helices in the core of the structure (gray sphere) highlighting the dynamic breathing motion of the HA protein.

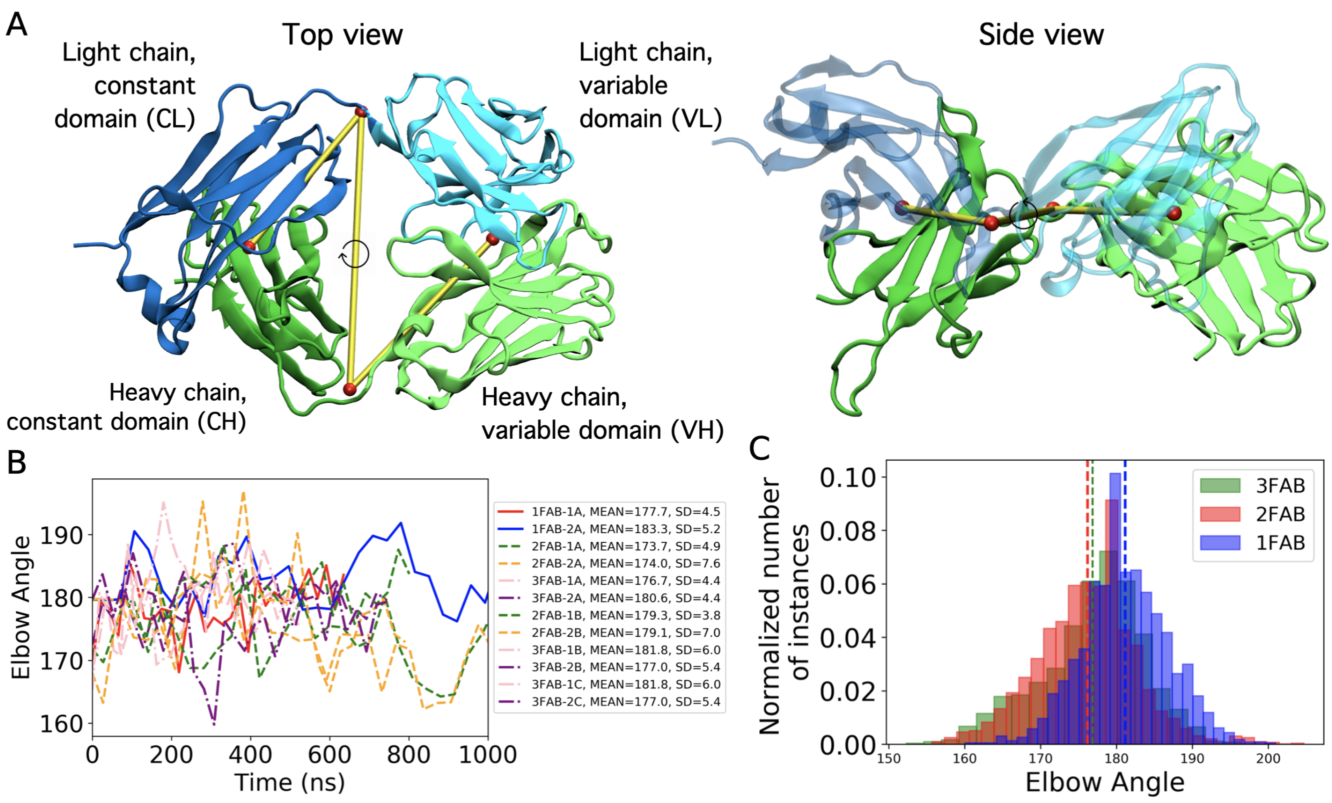


Figure S6: **The elbow angle of the HA-bound Fab domains.** (A) Illustration of the torsion angle used to calculate the elbow angle for the Fab domains, with constant domains shown in dark green (CH) and dark blue (CL), and variable domains shown in light green (VH) and light blue (VL). The elbow angle is defined as the torsion angle between the COMs of the variable domain, hinge region of the light chain (hl), constant domain, and hinge region of the heavy chain (hh). (B) Time series of the elbow angles computed from each Fab during simulations. (C) The histogram depicts the distribution of elbow angles, revealing differences between 1-Fab and multiple-Fab systems.

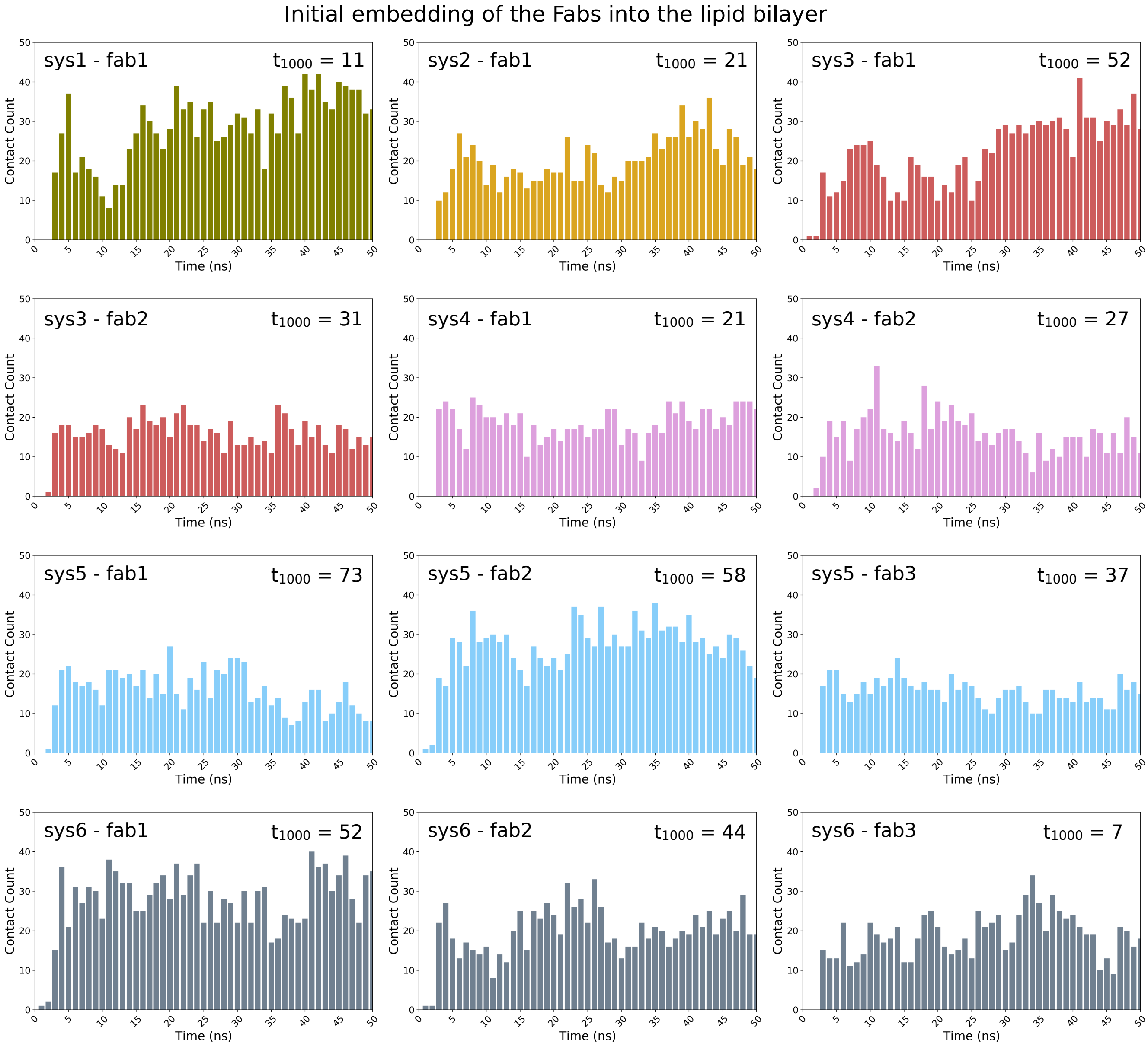


Figure S7: **The initial embedding of the Fabs into the membrane.** Plots show the number of contacts (see Methods for contact definition) between the Fabs and the lipid bilayer at each timestep for the first 50 ns after the end of the grid potential simulations where Fab domains were represented as repulsive potentials. Within a few nanoseconds, all Fabs embed into the membrane and remain stably bound until the end of the simulation ($t = 1\mu$s), for which we explicitly label the contact number in each plot.

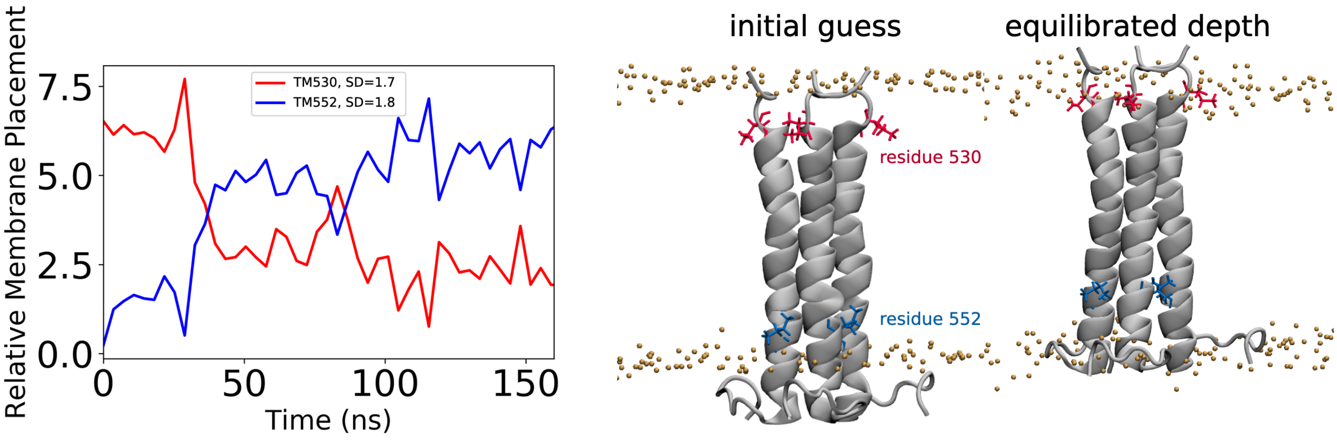


Figure S8: **Equilibration of the relative depth of the TM domain prior to complex assembly.** Convergence from the equilibrium simulations was tracked by observing the distance between the upper and lower phosphate layers of the membrane and COM of residues 530 (red) and 552 (blue) of the alpha helix of the TM domain.

Table S1: Nucleotide sequences (5′ to 3′) encoding wild-type, variable heavy chain and light chain of FISW84.

| | |
|---|---|
| Variable heavy chain | GAGGTGCAACTGCTTGAAAGCGGAGGCGGGCTGGTCCAGCCAGGCGGTTCCCTCAGACTTTCATGTGCCGCCAGCGGATTCACC<br>TTTTCTTCTTACGGGATGGCGTGGGTACGGCAGGCCCCCGGCAAAGGTTTGGAATGGGTCAGTTTCATTTCAGCAACTGGTTTG<br>AGCACATACTTCGCTGATTCCGTTAAGGGTCGCTTTACGATCAGTCGAGACACGACTAAAAATACCTTGTACCTCCAAATGAAT<br>AGTCTTCGGGCGGATGACACGGCTGTCTATTTTTGCGCACGAATGCGCAGAACTATGATAGCTTTTGGAGGGAATGACTTTTGG<br>GGTCAAGGCACATTGGTTACGGTGTCCTCC |
| Variable light chain | GAGGTGGTGATGACTCAGAGTCCCGCCACCCTGAGCGTGAGCCCAGGGGAGGGCGCCACACTGTCCTGCAGGGCTAGCCAGTCT<br>GTGAACACCAATGTGGCCTGGTATCAGCAGAAGCCAGGCCAGGCCCCCAGGCTGCTGATTTACGGCGCCTCTACCAGAGCCACC<br>GGCATCCCCGCTAGATTCAGCGGAAGCGGCTCCGGCACAGAGTTCACCCTGACCATTAGTACCCTGCAGTCCGAGGACTTTGCA<br>GTGTACTATTGCCAGCAGTACAGCAACTGGCCTCCTATTACCTTTGGCCAGGGAACAAGACTGGAAATCAAG |

Table S2: Nucleotide sequences of primers to generate mutations of FISW84.

| **Primer** | **Sequence (5′ to 3′)** |
|---|---|
| Heavy-K126A-F | CGGTGTCCTCCGCCTCTACCGCGGGACCAAGCGTTTTCCCATTG |
| Heavy-K126A-R | CAATGGGAAAACGCTTGGTCCCGCGGTAGAGGCGGAGGACACCG |
| Heavy-K126E-F | CGGTGTCCTCCGCCTCTACCGAGGGACCAAGCGTTTTCCCATTG |
| Heavy-K126E-R | CAATGGGAAAACGCTTGGTCCCTCGGTAGAGGCGGAGGACACCG |
| Heavy-K215A-F | CAATCACAAACCATCCAACACCGCAGTCGACAAAAGAGTCGAGC |
| Heavy-K215A-R | GCTCGACTCTTTTGTCGACTGCGGTGTTGGATGGTTTGTGATTG |
| Heavy-K215E-F | CAATCACAAACCATCCAACACCGAAGTCGACAAAAGAGTCGAGC |
| Heavy-K215E-R | GCTCGACTCTTTTGTCGACTTCGGTGTTGGATGGTTTGTGATTG |
| Heavy-N213A-F | CGTCAATCACAAACCATCCGCCACCAAAGTCGACAAAAGAGTCG |
| Heavy-N213A-R | CGACTCTTTTGTCGACTTTGGTGGCGGATGGTTTGTGATTGACG |
| Heavy-N213W-F | CGTCAATCACAAACCATCCTGGACCAAAGTCGACAAAAGAGTCG |
| Heavy-N213W-R | CGACTCTTTTGTCGACTTTGGTCCAGGATGGTTTGTGATTGACG |
| Heavy-Q13A-F | CTTGAAAGCGGAGGCGGGCTGGTCGCGCCAGGCGGTTCCCTCAG |
| Heavy-Q13A-R | CTGAGGGAACCGCCTGGCGCGACCAGCCCGCCTCCGCTTTCAAG |
| Heavy-Q13W-F | CTTGAAAGCGGAGGCGGGCTGGTCTGGCCAGGCGGTTCCCTCAG |
| Heavy-Q13W-R | CTGAGGGAACCGCCTGGCCAGACCAGCCCGCCTCCGCTTTCAAG |
| Heavy-R19A-F | CAGCCAGGCGGTTCCCTCGCACTTTCATGTGCCGCCAGCGG |
| Heavy-R19A-R | CCGCTGGCGGCACATGAAAGTGCGAGGGAACCGCCTGGCTG |
| Heavy-R19E-F | CAGCCAGGCGGTTCCCTCGAACTTTCATGTGCCGCCAGCGG |
| Heavy-R19E-R | CCGCTGGCGGCACATGAAAGTTCGAGGGAACCGCCTGGCTG |
| Heavy-S124A-F | GGTTACGGTGTCCTCCGCCGCTACCAAGGGACCAAGCGTTTTC |
| Heavy-S124A-R | GAAAACGCTTGGTCCCTTGGTAGCGGCGGAGGACACCGTAACC |
| Heavy-S124W-F | GGTTACGGTGTCCTCCGCCTGGACCAAGGGACCAAGCGTTTTC |
| Heavy-S124W-R | GAAAACGCTTGGTCCCTTGGTCCAGGCGGAGGACACCGTAACC |
| Heavy-T125A-F | GTTACGGTGTCCTCCGCCTCTGCCAAGGGACCAAGCGTTTTC |
| Heavy-T125A-R | GAAAACGCTTGGTCCCTTGGCAGAGGCGGAGGACACCGTAAC |
| Heavy-T125W-F | GTTACGGTGTCCTCCGCCTCTTGGAAGGGACCAAGCGTTTTC |
| Heavy-T125W-R | GAAAACGCTTGGTCCCTTCCAAGAGGCGGAGGACACCGTAAC |
| Light-K127A-F | CCCCTTCTGACGAGCAGCTGGCGTCTGGCACAGCCAGCGTGGTG |
| Light-K127A-R | CACCACGCTGGCTGTGCCAGACGCCAGCTGCTCGTCAGAAGGGG |
| Light-K127E-F | CCCCTTCTGACGAGCAGCTGGAGTCTGGCACAGCCAGCGTGGTG |
| Light-K127E-R | CACCACGCTGGCTGTGCCAGACTCCAGCTGCTCGTCAGAAGGGG |
| Light-K184A-F | CTAGCACCCTGACACTGAGCGCGGCCGACTACGAGAAGCACAAG |
| Light-K184A-R | CTTGTGCTTCTCGTAGTCGGCCGCGCTCAGTGTCAGGGTGCTAG |
| Light-K184E-F | CTAGCACCCTGACACTGAGCGAGGCCGACTACGAGAAGCACAAG |
| Light-K184E-R | CTTGTGCTTCTCGTAGTCGGCCTCGCTCAGTGTCAGGGTGCTAG |
